# Boosting bactericidal immunity of a recombinant *Mycobacterium smegmatis* strain via zinc-dependent ribosomal proteins

**DOI:** 10.1101/2023.12.11.571163

**Authors:** Shivani Singh, David Kanzin, Sarah Chavez, N. Alejandra Saavedra-Avila, Tony W. Ng, Regy Lukose, Oren Mayer, John Kim, Bing Chen, Mei Chen, Steven A. Porcelli, William R. Jacobs, Sangeeta Tiwari

## Abstract

Tuberculosis (TB) continues to be a major global health burden and kills over a million people annually. New immunization strategies are required for the development of an efficacious TB vaccine that can potentially induce sterilizing immunity. In this study, we first confirmed that various strains of the IKEPLUS vaccine confer a higher survival benefit than BCG in a murine model of intravenous *Mycobacterium tuberculosis* (Mtb) infection. We have shown that there was a significant increase in the expression of the Rv0282 when IKEPLUS was grown in low zinc and iron containing Sauton medium. We confirmed on biofilm assays that zinc plays a vital role in the growth and formation of *Mycobacterium smegmatis* (*M. smegmatis*) biofilms. IKEPLUS grown in low zinc media led to better protection of mice after intravenous challenge with very high dosage of Mtb. We also showed that various variants of IKEPLUS induced apoptotic cell-death of infected macrophages at a higher rate than wild type *M. smegmatis*. We next attempted to determine if zinc containing ribosomal proteins such as rpmb2 could contribute to protective efficacy against Mtb infection. Since BCG has an established role in anti-mycobacterial efficacy, we boosted BCG vaccinated mice with rmpb2 but this did not lead to an increment in the protection mediated by BCG.

## Introduction

Tuberculosis (TB) continues to be a major global health burden and kills over a million people annually (1). Despite its questionable efficacy to prevent adult pulmonary TB, the attenuated *Mycobacterium bovis* strain Bacille Calmette-Guerin (BCG) is the only licensed TB vaccine and continues to be widely used (2). By virtue of being a live vaccine, BCG has restricted use in subjects most vulnerable to TB such as individuals living with HIV; and the generation of safer and more efficacious vaccines is therefore paramount. This requires a better understanding of the mechanisms that *Mycobacterium tuberculosis* (Mtb) deploys to evade the host immune response. One such mechanism is known to involve the specialized mycobacterial type VII secretion system known as ESX-1 (3, 4). An intact *esx-3* locus has been shown to be essential for the growth and survival of Mtb (5, 6) but how exactly the *esx-3* locus contributes to mycobactericidal evasion is unknown. Expression of the *esx-3* genes is regulated by zinc, suggesting that zinc may have a key role to play in Mtb virulence (7) (8). We had previously constructed a vaccine strain IKEPLUS by introducing the Mtb *esx-3* genes into a *Mycobacterium smegmatis* (*M. smegmatis* /msmeg) Δesx-3 mutant strain (IKE=immune killing evasion) (9). Compared to BCG, the immunization of C57BL/6 mice with the IKEPLUS strain led to marked reductions in lung, spleen and liver bacterial burdens as well as prolonged survival after Mtb challenge (9). Inoculation with IKEPLUS induced a more robust Th1 cytokine milieu with a rapid IFN-γ and IL-12 response and increased phagosome maturation during macrophage infection.

We then sought to determine the mechanism of bacterial killing in IKEPLUS-immunized mice as a crucial next step. Since protective immunity induced by IKEPLUS was dependent on antigen - specific CD4 T cell responses, we hypothesized that the specificity of this response would be a critical feature of its protective phenotype. Using IFN-γ enzyme-linked immunosorbent spot (ELISPOT) assays on splenocytes from IKEPLUS-vaccinated mice, we identified an immunogenic peptide within the mycobacterial ribosomal protein RplJ (10). In a complementary approach, we generated major histocompatibility complex (MHC) class II-restricted T cell hybridomas from IKEPLUS-vaccinated mice and demonstrated that the CD4LJT cell response recognized several components of the mycobacterial ribosome and was localized in the lungs after Mtb challenge. We then extended our investigations by broadly screening all structural proteins of the Mtb ribosome for their ability to be targeted by CD4 T cells in IKEPLUS immunized mice and showed that the mycobacterial ribosome was highly immunogenic (11). Induction of strong multifunctional Th1 CD4 T cell responses and reduction of lung pathology and bacterial burdens was noted in RplJ-boosted mice after Mtb challenge (11).

With the above background, here we hypothesized that vaccination strategies directed at priming CD4LJT cells against zinc-dependent ribosomal proteins such as rpmb2 could be considered an approach to enhance vaccine efficacy against Mtb. Subdominant or cryptic antigens may have the potential to induce CD4+ T cell responses that can result in bactericidal and protective immunity. First, we illustrated a survival difference when mice were immunized with IKEPLUS strains prior to Mtb infection. Since secretory antigens such as ESAT-6 and CFP-10 are secreted when Mtb is grown in Sauton media but not in 7H9 media (12), we grew all msmeg strains in Sauton. It has also been reported that some of the genes in the ESX-3 region are induced by iron (13) and that ESX-3 is under the regulation of Zur-FurB regulon (7). We demonstrated a fivefold increase in the expression of the Rv0282 (first gene in the ESX-3 operon) when msmeg strains were cultured in either low iron or low zinc containing Sauton. Mice vaccinated with IKEPLUS/ mc^2^5009 grown in low zinc Sauton media were better protected after Mtb challenge. Finally, since rpmb2 is a zinc dependent ribosomal protein, we constructed rpmb2 specific T cell hybridomas from intravenous IKEPLUS vaccinated mice and tested their reactivity after incubating them with bone marrow derived dendritic cells that had been infected with H37Rv. Since BCG has an established protective efficacy against TB, we boosted BCG-vaccinated mice with subcutaneous rpmb2 but found that this did not have an incremental effect on the protective efficacy of BCG.

## Methods

### Bacterial strains, plasmids, phages and media

All msmeg strains used in this study were derived from the laboratory strain mc^2^155. Msmeg strains were cultured in 7H9 liquid medium (Difco, Becton-Dickinson) containing 0.5% glycerol, 0.2% dextrose and 0.05% Tween-80 for mutant construction. The strain was cultured in regular Sauton medium (RS) containing 0.05% Tween-80 for animal infections (9). Sauton medium used for the growth of Msmeg strains consisted of 60 ml/l of glycerol, 0.5 g/l of KH_2_PO_4_, 2.0 g/l of citric acid monohydrate, 4 g/l of asparagine, 0.5gm/l of magnesium sulphate, 1mg/l of zinc sulfate and 50mg/l of ferric ammonium citrate. pH of the solution was adjusted to 7:00-7.2 with the addition of 10N sodium hydroxide. For low zinc and low iron containing Sauton media, the final concentration of zinc was (100μg/l) and the final concentration of iron was 0.5mg/l. The BCG Danish SSI strain used for immunizations was from AERAS and the Mtb strain was an H37Rv derivative obtained from the Trudeau Institute. Mtb and BCG were cultured using Middlebrook 7H9 liquid medium (Difco, Becton-Dickinson) containing 10% OADC (oleic acid/albumin/dextrose/catalase), 0.5% glycerol and 0.05% Tween-80. Media were supplemented with ‘as required’ antibiotics hygromycin (50μg/mL) or apramycin (20μg/mL). Colony counts were plated on 7H10 agar plates containing 10% OADC, 0.5% glycerol and 0.05% Tween-80. Descriptions of all strains and plasmids used in this study is provided in Supplementary Tables 1, 2 and 3.

### Generation of IKEPLUS

Specialized transducing phage (phAE543), as described previously (9), (14), was used to generate MsΔ*esx-3* (Ms0615-Ms0626) or mc^2^6462. Transductions were plated on 7H10 plates containing 150 µg/ml hygromycin. Hygromycin resistant clones were screened for deletion by polymerase chain reaction (PCR) and further confirmed by whole genome sequencing. Antibiotic marker was removed using phage pHAE280 to generate the unmarked strain (mc^2^6463 or IKE) followed by transformation of this strain with cosmid pYUB2098 (Rv0278- Rv303)::kan to generate mc^2^6456. As this strain contains an integrase that can lead to instability of the cosmid pYUB2098(Rv0278-Rv303)::kan, mc^2^6456 was transformed with a plasmid bearing resolvase (pYUB2099) and named mc^2^7159 (Stable IKEPLUS). Furthermore, in order to transform this strain with an immunoreporter without incorporating an antibiotic marker, we deleted *leuCD* gene in mc^2^7159 to generate mc^2^7170 and unmarked it to make mc^2^7173 using pHAE280. Finally, mc^2^7173 was transformed with immunoreporter *PBRL635* containing immunogenic epitopes of ESAT-SINFIKEL-OVAII on a plasmid with leuCD genes as a selection marker (instead of antibiotic) and called mc^2^7257 (SIPΔleuCD::pBRL635leuCD). In our murine survival experiments, we compared this IKEPLUS strain with the previously generated IKEPLUS strain mc^2^5009 (9).

### Q-PCR

In to determine if the expression of ESX-3 in IKEPLUS changes in response to low iron and low zinc conditions, we performed Q-PCR. mc^2^5009 and mc^2^7182 were cultured in regular Sauton (RS), low iron Sauton (LFeS) and low zinc Sauton (LZnS) media. Cultures were pelleted and suspended in “RNA protect” for storage until further processing. For the purpose of isolating RNA, samples were centrifuged and the pellets were suspended in 1ml trizol, followed by bead-beating (3times/30sec). Supernatants were collected and an equal volume of ethanol was added. Further purification was performed as per manufacturers’ protocol (Zymo-spin RNA purification kit, Zymo Research Corporation). c-DNA was synthesized as per protocol in the Superscript III Reverse Transcriptase kit (Invitrogen#18989-093). This was followed by Q-PCR using Sybr green mix (Promega). 40 cycles were performed and relative expression of Rv0282 was determined in RS, LFeS and LZnS media after normalizing with 16S RNA.

### Murine infections

C57BL/6 mice (6-8 weeks old) were obtained from the Jackson Laboratory. All msmeg strains used for immunization of mice were cultured either in RS or LZnS media. BCG Danish SSI was grown in 7H9 media. Before inoculation, bacteria were washed twice with PBS plus 0.05% Tween 80. Subsequently, bacteria were sonicated twice for 10 seconds each time, and then diluted to achieve the desired densities. Different groups of mice were immunized with PBS intravenously (i.v.) or BCG/ IKEPLUS strains grown in either RS or LZnS media. After 7-8-weeks of immunization mice were challenged with high dose of Mtb strain H37Rv (5.5 x10^7^ or 10^8^ CFU/mice) via the i.v. route. The mice were observed without any further intervention for survival studies. Statistical analysis was done using log rank test. The animal protocol was approved by the University of Texas at El Paso and Albert Einstein College of Medicine Institutional Animal Care and Use Committee (IACUC) protocols, as per NIH principles for animal usage.

### Boosting BCG-primed C57BL/6 mice with rpmb2 ribosomal protein

C57BL/6 mice were vaccinated with 5 x10^6^ CFU’s of BCG Danish via the SQ route. After an interval of 6 weeks and 8 weeks, the BCG primed mice were boosted via the subcutaneous route with 25μg of rpmb2 mixed with the adjuvant CpG ODN type B. CpG oligodeoxynucleotides (or CpG ODN) are short single-stranded synthetic DNA molecules that contain a cytosine triphosphate deoxynucleotide (C) followed by a guanine triphosphate deoxynucleotide (G). They are a new class of Th1-type immune stimulants that stimulate the TLR9 receptor and in doing so have proven to be a highly efficacious adjuvant for mucosal vaccines against infectious diseases. The prime-boosted mice were aerosol infected with H37Rv at 12 weeks after BCG prime and the mice were euthanized for organ retrieval (lung, spleen and thoracic lymph nodes) and CFU enumeration at 10 weeks after H37Rv challenge.

### Polycaspase activation assays

Bone marrow macrophages (BMM) were isolated and differentiated using L929 supernatants as described previously (14). Peritoneal macrophages (PM) were prepared by injecting 2ml of 3% Brewer Thioglycolate into the peritoneal cavity of mice. Four days after inoculation, mice were euthanized and peritoneal macrophages were extracted from peritoneal cavity by injecting 5ml of

PBS and withdrawing the same volume of fluid from the peritoneal cavity. The cells were centrifuged, washed and suspended in complete DMEM medium for plating. After 24h cells were washed to removed suspended cells and adhered macrophages were cultured. After 6-7 days macrophages were trypsinized, counted and seeded for infection assays. All the smegmatis strains were grown in LZnS media. 24 hours after seeding, macrophages were infected for 3h with the respective strains at an MOI of 1:10. Cells were washed and stained with FLICA 660 PolyCaspase kit (Immunochemistry technologies Catalog# 9120) 24 or 48h post-infection to measure apoptosis as a surrogate measure of active caspase enzymes. For flow cytometry, cells were stained as per manufacturer’s instructions and samples were read on the FACS Aria flow cytometer.

### Biofilm Assay

Sauton Media was prepared as mentioned above. Zinc and iron supplements were added as indicated. Log phase cultures of the desired strains were pelleted by centrifugation (4000rpm/10min/RT). This was followed by resuspending the cells in 10ml Sauton media and re-suspending the centrifuged pellet with Sauton media to the original volume of the culture. Biofilm growth was performed in a 12-well polystyrene plate. Wells were inoculated with cells at a ratio of 1:50 cells to the total volume of the media in the well. Plates were wrapped in the foil and incubated at 37°C, 5% CO2 and checked weekly for the biofilm growth.

### Expression of Mtb ribosomal protein rpmb2 in *E. Coli* plasmids

Production of recombinant Mtb ribosomal protein rpmb2 was undertaken in collaboration with the protein core using *E. coli* harboring plasmids that encoded these ribosomal proteins. Auto-induction was done using Isopropyl β-D-1-thiogalactopyranoside (IPTG) as the auto-induction reagent. Purification was undertaken using with nickel affinity chromatography, size-exclusion chromatography and ion exchange chromatography, followed by overnight dialysis against cell-culture grade PBS. Next, endotoxin was reduced by overnight incubation with endotoxin removal resin. Finally, protein concentration was measured by UV absorbance at 280nm and endotoxin load was determined using a Limulus amebocyte lysate assay.

### Generation of rpmb2 specific T cell hybridoma and IL-2 ELISA

C57BL/6 mice were immunized with 10×10^6^ CFU of IKEPLUS via the intravenous route. After an interval of 4 weeks and 6 weeks, these mice were injected with 25μg rpmb2 mixed with 20μg of CPG ODN via the intraperitoneal route. 2 weeks after the last boost, the mice were sacrificed and spleen and mesenteric lymph nodes were harvested. Using the Miltenyi Biotec kit (catalogue #130-104-454), that used CD4+ T cell biotin-antibody cocktail and anti-biotin microbeads, CD4 T cells were isolated from the spleen and mesenteric lymph nodes and incubated with rpmb2 and bone marrow derived dendritic cells for 3 days. Next, these CD4 T cells were fused with thymoma cells as per previously published guidelines (15, 16). The fused cells (hybridomas) were expanded and were then subjected to a limiting dilution. The primary hybridomas were pre-screened for TCR-β expression using flow cytometry. BCG infected bone marrow derived dendritic cells were incubated overnight with rpmb2 and Ag85B specific T cell hybridomas. Antigen specificity was confirmed with an IL-2 ELISA assay. Supernatants were harvested and the levels of IL-2 were quantitated by a capture ELISA as previously described (17). Absorbance values were determined at 450 nm on a Wallace 1420 Victor2 microplate reader (PerkinElmer, Waltham, MA) and values for IL-2 were determined by using a standard curve generated by using known concentrations of purified recombinant murine IL-2 (BD Biosciences).

## Results

### Immunization with IKEPLUS strains protect mice against challenge with high dose Mtb

6-8 weeks old C57BL/6 mice (n=10) were sham immunized with i.v. PBS, SQ BCG (10^7^ CFU/mouse) and SQ IKEPLUS strains SIPΔleuCD::pBRL635leuCD (mc^2^ 7257) or IKEPLUS (mc^2^ 5009) with approximately 5×10^8^ CFU/mouse grown in the RS media as per the strategy illustrated in Fig.1A. After 8-weeks of immunization, mice were challenged with high dose of i.v. Mtb strain H37Rv (5.5 x10^7^ CFU/mouse). The mice were observed for survival studies as shown in Fig.1B. Differences in the survival curves were significant for PBS versus SIPΔleuCD::pBRL635leuCD strain (p=0.0001, log rank test) as well as PBS versus IKEPLUS strain (p=0.0001, log rank test). The difference observed between the protective efficacies of the two IKEPLUS strains was insignificant and could be attributed due to the difference in the CFU delivery to the mouse lung (target dose was 5×10^8^ /mouse but CFU delivered was 7×10^8^ for mc^2^5009 strain). In addition, there was a survival benefit when mice were immunized with SQ BCG but the statistical significance was much smaller compared to PBS versus IKEPLUS strains (p=0.01).

**Figure 1:**
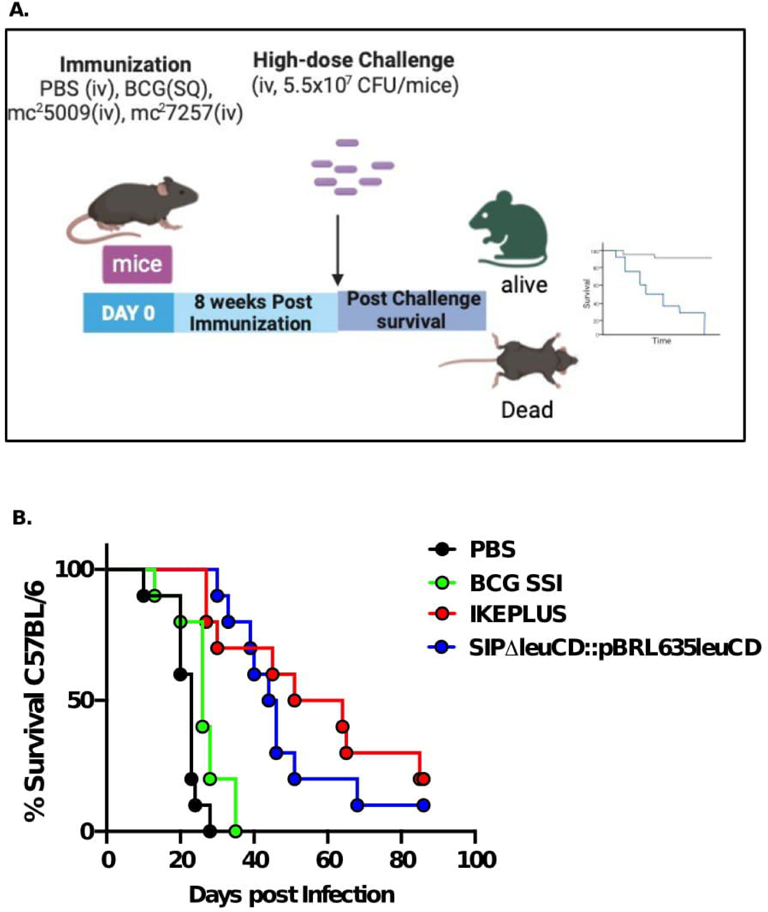
Immunization with IKEPLUS strains protect mice against challenge with high dose Mtb. Figure lA: 6-8 weeks old C57BL/6 mice (n=lO) were sha1n immunized with i.v. PBS, SQ BCG (10^7^ CFU/mouse) and SQ IKEPLUS strains SIP leuCD::pBRL635leuCD (mc^2^ 7257) or IKEPLUS (111c^2^ 5009) with approxi1nately 5×10^8^ CFU/1nouse grown in tl1e RS media. After 8-weeks of <u>immunization, mice</u> were challenged with high dose of i.v. Mtb strain H37Rv. **Figure.lB.** Differences in the survival curves were significant for PBS versus SIP leuCD::pBRL635leuCD strain (p=0.0001, log rank test) as well as PBS versus IKEPLUS strain (p=0.0001, log rank test). In additio11, there was a survival benefit when mice were i1nmunized witl1 SQ BCG but the statistical significance was mucl1 smaller co111pared to PBS versus IKEPLUS strains (p=0.01).

### LZnS medium enhances IKEPLUS mediated protection against high dose of i.v. Mtb challenge

Here, we tested whether modulating the level of zinc and iron in Sauton growth medium can regulate the expression of the Mtb ESX-3 gene expressed in *M. smegmatis*. We found that there was a fivefold increase in the expression of the Rv0282 (first gene in the ESX-3 operon) when IKEPLUS was grown in LZnS and LFeS compared to RS medium (Fig. 2A). Biofilm assays showed that wild type *M. smegmatis* mc^2^155 grown in 7H9 media formed thin biofilms but the same strain grown in Sauton medium formed thick biofilms (Fig. S1A). Additionally, when old Sauton medium was supplemented with zinc and iron, *M. smegmatis* growth was augmented. **Figure S1B.** Next, we investigated the role of zinc on the growth of wild type *M. smegmatis*, IKEPLUS and IKE. Zinc concentrations were titrated to support the growth of *M. smegmatis.* We found that zinc plays a vital role in the growth and formation of *M. smegmatis* biofilms, and 100ug/l of zinc sufficiently supported the growth of *M. smegmatis* (Fig. S1B). Therefore, we choose low zinc (100ug/l) Sauton media in further infection studies. C57BL/6 mice (n=14) were immunized with i.v. PBS, SQ BCG (10^7^ CFU/mouse) and the IKEPLUS strain mc^2^5009 (5×10^7^/mouse) grown in RS or LZnS media. Seven weeks after immunization, mice were challenged with high dose of i.v. Mtb strain H37Rv (1 x10^8^ CFU/mouse). Mice were observed for survival studies. IKEPLUS/ mc^2^5009 grown in LZnS media led to better protection of mice after i.v. challenge with very high dosage of Mtb (1×10^8^/mouse) compared to mice immunized with IKEPLUS/ mc^2^5009 grown in RS (Fig. 2B, p=0.03). Differences in the survival curves were significant for both mc^2^5009/IKEPLUS grown in RS (p=0.001, log rank test) and LZnS (p=0.001, log rank test), when compared to IKEPLUS grown in PBS.

**Figure 2:**
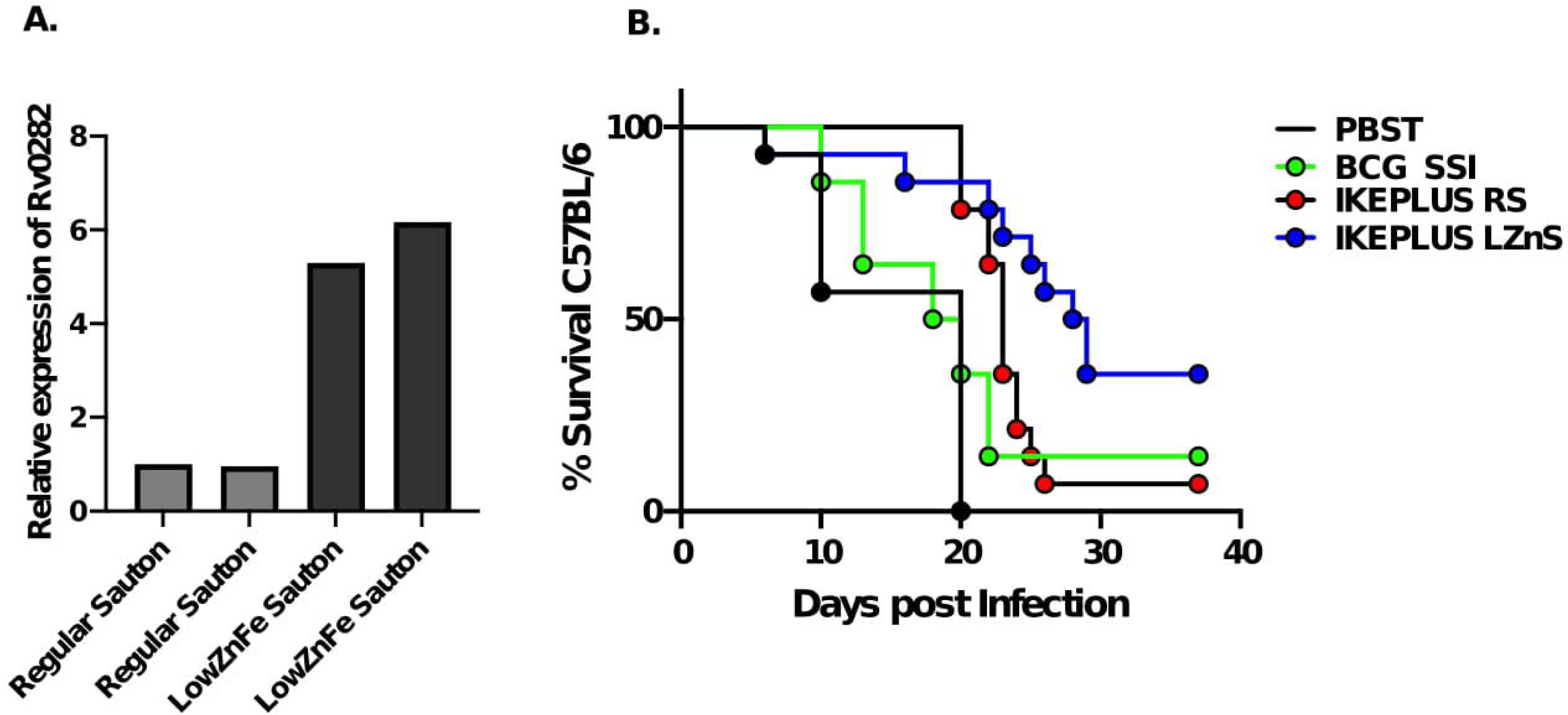
LZnS medium enhances IKEPLUS mediated protection against high dose of i.v. Mtb challenge. Figure 2A: A fivefold increase i11 the expressio11 of the Rv0282 (first gene in the ESX-3 operon) wl1en IKEPLUS was grown i11 LZnS and LFeS co1npared to RS 1nedium. Figure 2B: C57BL/6 111ice (n=l4) were immunized with i.v. PBS, SQ BCG (10^7^ CFU/mouse) a11d the IKEPLUS strain mc^2^5009 (5×l0^7^/mouse) grown in RS or LZnS media. Seven weeks after immunization, mice were challenged with higl1 dose of i.v. Mtb st1·ain H37Rv (1 xl0^8^ CFU/mouse). IKEPLUS/ mc^2^5009 grow11 in LZnS media led to better protectio11 of mice after i.v. challenge with very high dosage of Mtb (lxl0^8^/mouse) compared to mice immunized with IKEPLUS/ mc^2^5009 grown in RS (p=0.03). Differences in the survival curves were significant for both mc^2^5009/IKEPLUS grown in RS (p=0.001, log rank test) and LZnS (p=0.001, log rank test), when compared to IKEPLUS grown i11 PBS.

### IKEPLUS induces apoptotic cell-death of infected cells macrophages

Bone marrow macrophages (BMM) and peritoneal macrophages (PM) were infected with *M. smegmatis* strains grown in LZnS media at an MOI of 1:10 for 3h. After 24 or 48h post-infection, macrophages were washed and stained with FLICA (660 Poly Caspase kit) that detects active caspase enzymes involved in apoptosis. Stained macrophages were read on FACSAria flow cytometer. For negative control, we used uninfected (UI) macrophages and for positive control macrophages were stimulated with either LPS (100ng/ml) in combination with IFNγ (200U/ml) or Staurosporine (1μM overnight) for 24 or 48hours. We noted a 3-4 fold increase in the apoptosis rate of BMM or PM macrophages treated with LPS (100ng/ml) in combination with IFNγ (200U/ml) or Staurosporine (1μM overnight); as compared to uninfected macrophages (Fig.3A). There was an approximately 20% increase in the apoptosis observed in BMM treated with various variants of IKEPLUS compared to wild type mc^2^155 within 24 hours (Fig. 3B). This increment decreased to 10% but was still at 48h (Fig.3B). Infected PMs did not illustrate a significant difference in the activation of caspase/apoptosis at 24h but a 30% increase in apoptosis was noted in the samples infected with IKEPLUS variants compared to wildtype mc^2^155 at 48h (Fig. 3C).

**Figure 3:**
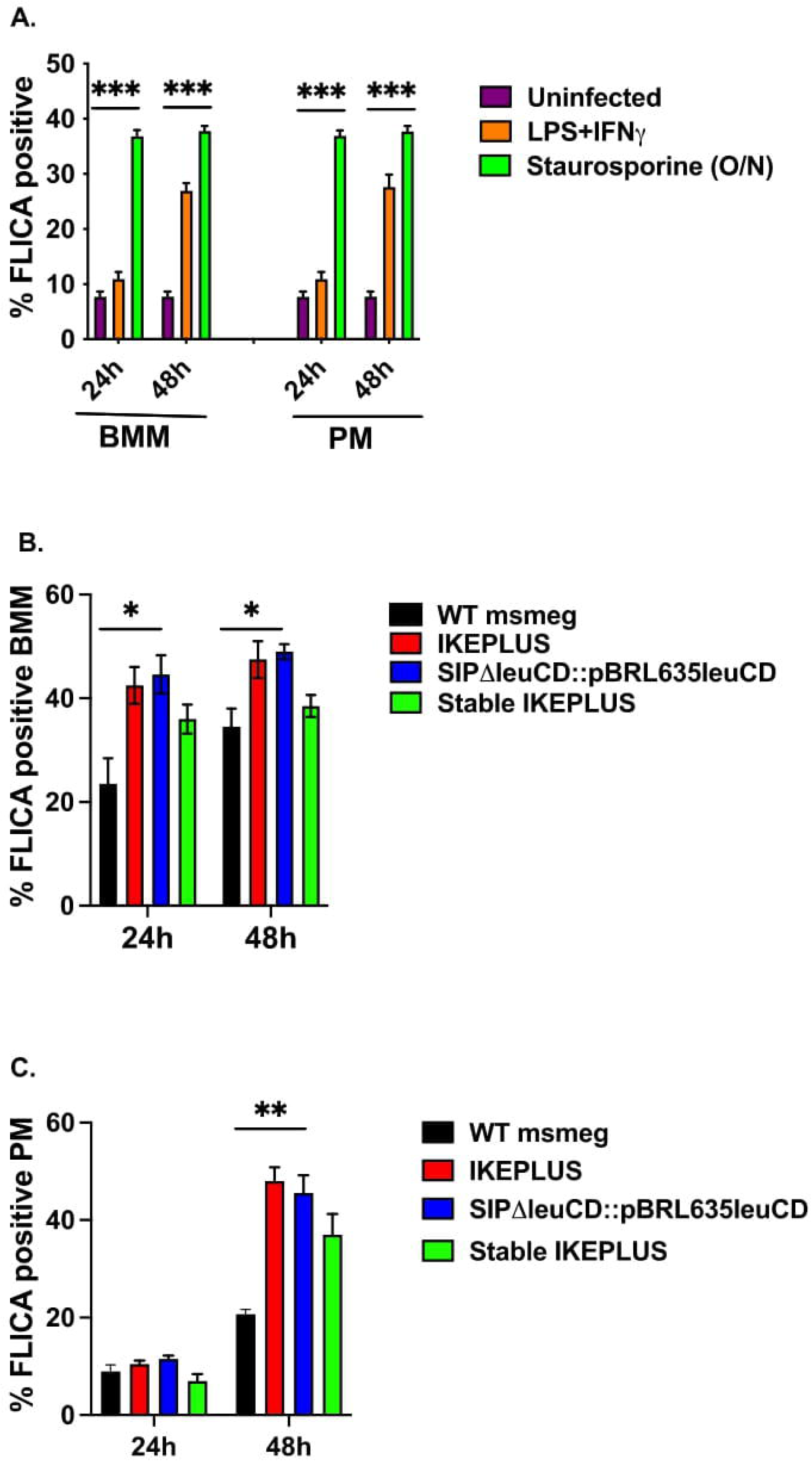
IKEPLUS induces apoptotic cell-death of infected cells macrophages. Figure 3A: A 3-4 fold increase in the apoptosis rate of bone marrow macrophages (BMM) or pe1·itoneal 1nacrophages (PM) treated with LPS (lOOng/ml) in combination with IFNy (200U/1nl) or Staurosporine (lµM overnight); as compared to uninfected macrophages was noted. Figure 3B: There was an approxi1nately 20% increase in the apoptosis observed in BMM treated with various variants of IKEPLUS compared to wild type mc^2^155 witltin 24 hours. Tltis incre1nent decreased to 10% but was still at 48h. Figure 3C: Infected PMs did not illustrate a significant difference in the activation of caspase/apoptosis at 24h but a 30% increase in apoptosis was noted in the samples infected with IKEPLUS variants compared to wildtype mc^2^155 at 48h.

### rpmb2 does not augment BCG mediated protection against Mtb

Finally, we show an incremental increase in the secretion of IL-2 upon incubation of rpmb2 (zinc-dependent ribosomal protein) specific T cell hybridomas and Ag85B specific T cell hybridomas with H37Rv-infected bone marrow derived dendritic cells (BMDC) (Fig. 4A-C). IL-2 secretion increased with an increase in the MOI of H37Rv infection of BMDC’s and was positive upon incubation with the ribosomal protein rpmb2. Ag85B specific T cell hybridomas were used as a positive control and did not stimulate IL-2 secretion upon incubation with rpmb2.

**Figure 4:**
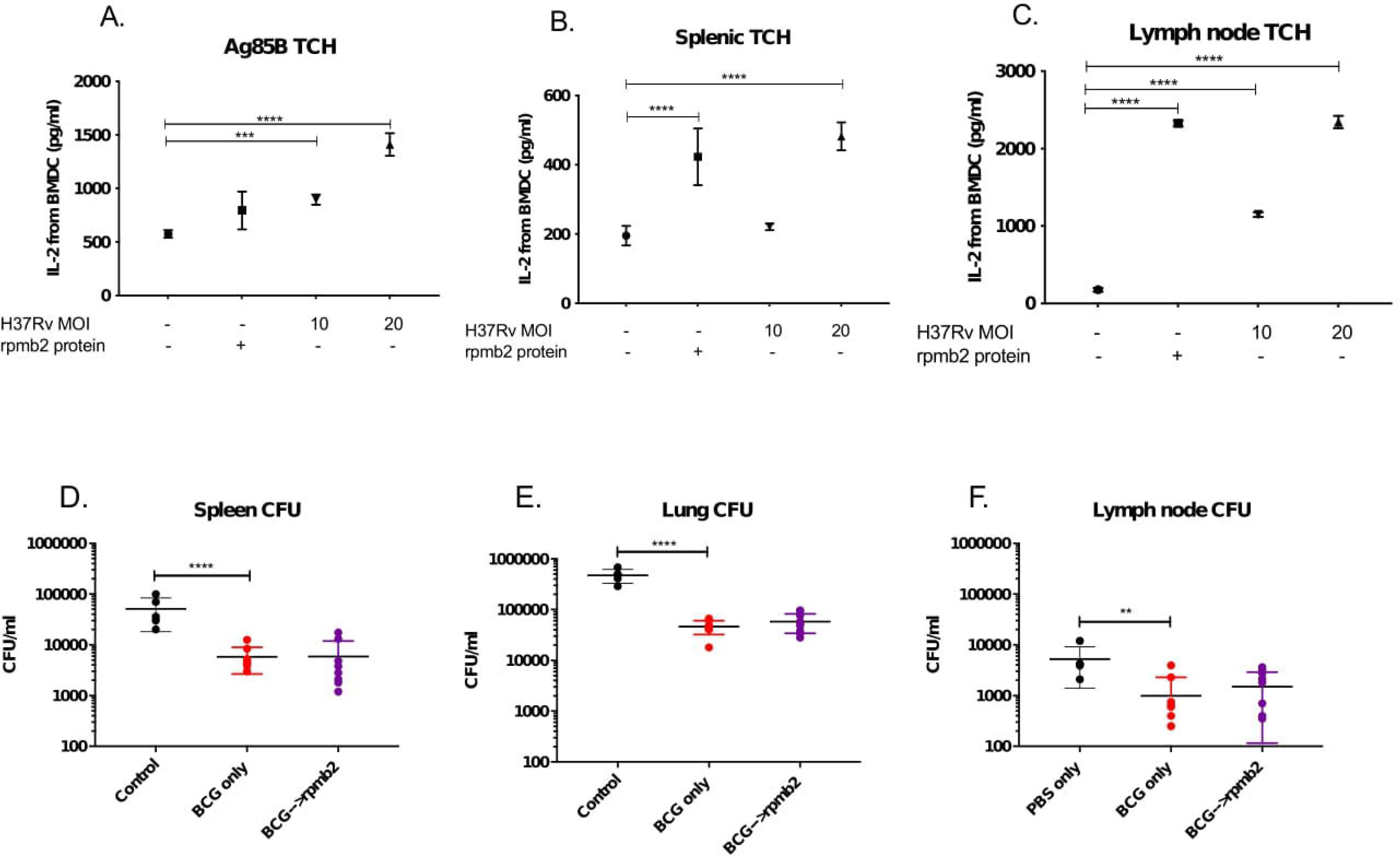
rpmb2 does not augment BCG mediated protection against Mtb. Figure 4A-C: An incremental increase in the secretion of IL-2 upon incubation of rpmb2 (zinc-dependent ribosomal protein) specific T cell hybridomas and Ag85B specific T cell hybridomas with H37Rv-infected bo11e 1narrow derived dendritic cells (BMDC) was noted. IL-2 secretion increased with an inc1·ease in the MOI of H37Rv infection of BMDC’s and was positive upon incubation with the ribosomal protein rpmb2 (4B, 4C). Ag85B specific T cell hyb1·idomas were used as a positive co11trol a11d did not stimulate IL-2 secretio11 upo11 incubatio11 with rp1nb2 (4A). Figure 4D-F. BCG vaccinated <u>mlce</u> were boosted witl1 rmpb2 but this did not lead to an increment i11 the protectio11 mediated by BCG, as illustrated by splee11, lu11g, and lyinph node CFUs (Fig. 4D-F).

Having established that rpm2-specific T cell hybridomas are stimulated by H37Rv-infected bone marrow derived dendritic cells, we next attempted to determine if rpmb2 could contribute to protective efficacy against Mtb infection in a mouse model of TB infection. BCG has an established role in anti-mycobacterial efficacy. We boosted BCG vaccinated mice with rmpb2 but this did not lead to an increment in the protection mediated by BCG, as illustrated by spleen, lung, and lymph node CFUs (Fig. 4D-F).

## Discussion

New immunization strategies are required for the development of an efficacious TB vaccine that can potentially induce sterilizing immunity and modulate the host immune system to eliminate virulent mycobacteria. In this study, we first confirmed our previous finding (9) that various strains of IKEPLUS confer a higher survival benefit than BCG. This may in part be due to the fact that BCG has a better protective profile against Mtb infection via the pulmonary route rather than the i.v. route that was used in our study (18) (19) (20). Next, we have shown that there was a fivefold increase in the expression of the Rv0282 (first gene in the ESX-3 operon) when IKEPLUS was grown in LZnS and LFeS medium compared to RS medium. This was confirmed on biofilm assays that showed that zinc plays a vital role in the growth and formation of *M. smegmatis* biofilms and IKEPLUS/ mc^2^5009 grown in LZnS media led to better protection of mice after i.v. challenge with very high dosage of Mtb. We surmise that this may be due to different pool of antigens induced /expressed under low zinc conditions such as ribosomal proteins (21). We also showed that various variants of IKEPLUS induced apoptotic cell-death of infected macrophages at a higher rate than wild type mc^2^155. Upon incubation of rpmb2 (zinc-dependent ribosomal protein) specific T cell hybridomas and Ag85B specific T cell hybridomas with H37Rv-infected bone marrow derived dendritic cells, IL-2 secretion increased with an increase in the MOI of H37Rv infection of BMDC’s. Having established that rpm2-specific T cell hybridomas are stimulated by H37Rv-infected bone marrow derived dendritic cells, we next attempted to determine if rpmb2 could contribute to protective efficacy against Mtb infection in a mouse model of TB infection. Since BCG has an established role in anti-mycobacterial efficacy, we boosted BCG vaccinated mice with rmpb2 but this did not lead to an increment in the protection mediated by BCG.

Candidate vaccines against Mtb have largely focused on targeting immunodominant antigens such as the Ag85 complex, ESAT-6 and TB10.4, to name a few (22), but enhanced protection has been elusive and safety issues have been frequent. Progress in this field has been hindered by a lack of knowledge about immune correlates of protection. The discovery that several known T cell epitopes of Mtb are derived from hyper-conserved T cell epitopes suggests that the recognition of these epitopes by the host may be beneficial to Mtb, thereby raising the possibility that there may be decoy antigens continued replication and transmission (23). Such CD4LJT cell epitopes, by virtue of not being directly involved in the host-pathogen coexistence, could represent more effective vaccine targets (10). Using a synthetic peptide library consisting of 880 non-overlapping 15-mers representing predicted I-Ab-presented epitopes, our group had previously shown that the specificity of the CD4LJT cells evoked by IKEPLUS and cross-reactive with Mtb were specific for structural proteins of the mycobacterial ribosome (10). This CD4LJT cell response was MHC class II restricted and dominated by the production of IFN-γ, suggesting the development of a Th1-like response. Taken together, these studies raise the possibility that immunization against mycobacterial ribosomal components primes the host immune response to recognize ribosomal epitopes when challenged with the virulent, ribosome-limited Mtb and that ribosome-specific CD4LJT cells are recruited to and expand in the lungs of IKEPLUS primed and Mtb-challenged mice (10). Ribosome vaccines are immunogenic, less toxic and better defined than whole-cell vaccines and studies in the 1960’ and 1970’s demonstrated the potential of ribosomal components as protective vaccines (24) (25). Components of the ribosome itself can act as adjuvants and stimulate both cell and humoral mediated immunity. But the mechanism of recognition, processing, and presentation of ribosomes following infection or immunization remains unclear, as do the underpinnings of the development of ribosome-specific T cell responses following infection (10).

One limitation of our study was that the bactericidal effects were most apparent when we administered IKEPLUS intravenously. This route is not the most feasible route for clinical trials. Thus, further improvements will be needed to optimize the efficacy of IKEPLUS vaccination for implementation in humans. This may involve prime-boost regimens. Taken together, our findings suggest that the incorporation of cryptic antigens into vaccination strategies may be a useful in promoting a protective CD4LJT cell response against Mtb. Others have shown that cryptic antigens are able to prime a CD4LJT cell response with greater TNF-α and IL-2 production and a higher proliferative capacity compared to immunodominant antigens (26). An Mtb genome-wide screen to identify targets of the major histocompatibility complex (MHC) class II-restricted CD4LJT cell responses highlighted the sheer breadth of epitopes recognized during Mtb infection, emphasizing the potential importance of designing multiepitope vaccines (27).

## Acknowledgements

NIH SC1GM140968 and BBRC (ST); R24AI134650-07A1, R37AI026170-37A1 (WRJ); R01 AI137344, AI127711, AI045889 (SAP);

---

**Supplementary Figure 1:**
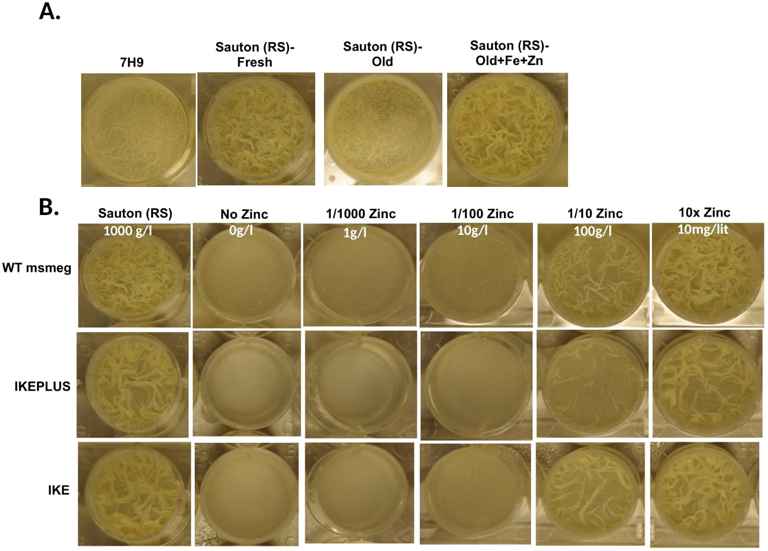
Zinc and Iron affect the growth of *M. smegmatis* in Sauton medium. Figure SlA: Biofilm assays showed that wild type *M smegmatis* mc^2^155 grown in 7H9 media formed thin biofilms but the same strain grown in Sauton medi1un formed thick biofilms. Additionally, whe11 old Sauto11 medium was suppleme11ted with zinc and iron, *M smegmatis* growth was augn1ented. **Figure SlB.** Next, we investigated the role of zinc on the growth of wild type *M smegmatis,* IKEPLUS and IKEPLUS. Zinc concentrations were titrated to s11pport the growth of *M smegmatis.* We found that zinc plays a vital role in the growth and formation of *M smegn1atis* biofilms, and 1OOug/1of zinc sufficiently supported the growth of *M smegmatis*.

