## Supplementary Tables 1, 2, 3 for "Boosting bactericidal immunity of a recombinant *Mycobacterium smegmatis* strain via zinc-dependent ribosomal proteins": Supplementary tables_Boosting bactericidal immunity of a recombinant Mycobacterium smegmatis strain via zinc-dependent ribosomal proteins.docx

| **Strain No** | **Strain name** | **Genotype** | **Description** | **Source/Reference** |
| --- | --- | --- | --- | --- |
| H37Rv | WT | Parent strain |  | Trudeau Institute |
| BCG SSI |  | Parent strain |  | AERAS |
| mc^2^155 | *ept-1* | Parent strain |  | (1) |
| mc^2^6462 |  | *ept-1* ∆(*Ms 0615-Ms0626*)*:: hyg*^R^, *suc*^S^ | Specialized transduction of mc^2^155 with phAE543 | Tiwari et al, (unpublished) |
| mc^2^6463 | IKE | *ept-1* ∆(*Ms 0615-Ms0626*) | Specialized transduction of mc^2^6462 with phAE280 | Tiwari et al, (unpublished) |
| mc^2^6456 | IKEPLUS_2 | *ept-1* ∆(*Ms 0615-Ms0626*)::  *attB_L5_* pYUB2098 *kan*^R^ | Transformation of mc^2^6463 with pYUB2098 (kan) | This study |
| mc^2^7159 | SIP | *ept-1* ∆(*Ms 0615-Ms0626*)::  *attB_L5_* pYUB2098∆kan∆integrase | Transformation of mc^2^6456 with pYUB2099 (containing delta gamma resolvase) | This study |
| mc^2^7170 |  | *ept-1* ∆(*Ms 0615-Ms0626*), *attB_L5_::pYUB2098*∆Kan*,* ∆(*leuCD,Ms2387-88*)*, :: hyg*^R^, *suc*^S^ | Specialized transduction of mc^2^7159 with pHAE763 | This study |
| mc^2^7173 |  | *ept-1* ∆(*Ms 0615-Ms0626*), *attB_L5_::pYUB2098*∆Kan*,* ∆(*leuCD,Ms2387-88*) | Specialized transduction of mc^2^7173 with pHAE280 | This study |
| mc^2^7257 | SIP∆*leuCD:: PBRL635* | *ept-1* ∆(*Ms 0615-Ms0626*), *attB_L5_::pYUB2098*∆Kan*,* ∆*leuCD + PBRL635 (leuCD)* | Transformation of mc^2^7173 with pBRL635 | This study |
| mc^2^5009 | IKEPLUS_1 |  |  | (2) |

**Table S1: List of strains used in the current study.** Abbreviations used *hyg^R^*: hygromycin cassette, *sacB*: counter selectable marker, k*an*^R^ kanamycin resistance.

**Table S2: List of plasmids and phasmids used in the current study:** Abbreviations used *hyg*^R^: hygromycin resistance, *suc*^S^: Sucrose sensitive, *kan*^R:^ Kanamycin resistance.

| **Plasmids/phages** | **Description** | **Reference** |
| --- | --- | --- |
| pYUB1136 | *attP_L5_λcosColE1 apr bla* | (2) |
| pYUB1336 | pYUB1136: :(Rv0278-Rv303) | (2) |
| pYUB2098 | pYUB1336(Rv0278-Rv303)::kan | This study |
| pYUB2099 | Gamma delta Resolvase, Hyg, SacB (to remove integrase) | This study |
| pBRL635 | Integrative plasmid with LeuCD | Tiwari et al. (unpublished) |
| phAE159 | Conditionally replicating shuttle phasmid vector | (2) |
| phAE543 | phAE159::pYUB1432 | (2) |
| phAE763 | ph159 with pYUB1572 | Tiwari et al. (unpublished) |

**Table S3: Primers used in the current study.**

| **Name of Primer** | **Sequence** | **Purpose** |
| --- | --- | --- |
| IKEPLUS_F setI (12907-13966) | GCCTCGACAGTTAGCTTATGCAATG | To confirm complementation |
| IKEPLUS_R setI  (12907-13966) | AACTCGGCGAGTTGGAGTTCG | To confirm complementation |
| IKEPLUS_F setII (22944-23441) | ACCGCACGACAGCAAGTAAC | To confirm complementation |
| IKEPLUS_R setII (22944-23441) | CCAAACCGACACCAAGAATCGG | To confirm complementation |
| Primers for Q-PCR | | |
| 16S-RT-F-Mo | GCC GTA AAC GGT GGG TAC TA | Primers for Q-PCR |
| 16S-RT-R-Mo | TGC ATG TCA AAC CCA GGT AA | Primers for Q-PCR |
| Rv0282-RT-F-Mo | ATT TCC ACC TCG CGT ATG CC | Primers for Q-PCR |
| Rv0282-RT-R-Mo | TGA GCA GCT TCA CGA CAT CC | Primers for Q-PCR |
